# Single-Platelet Mapping of Jugular, Puncture-Wound Thrombi Reveals the Spatial Evolution of Platelet Activation

**DOI:** 10.1101/2024.07.07.602390

**Authors:** Sung Rhee, Irina D. Pokrovskaya, Kelly Ball, Michael W. Webb, Jeffrey A. Kamykowski, Oliver Zhao, Elizabeth R. Driehaus, Maria A. Aronova, Sidney W. Whiteheart, Richard D. Leapman, Brian Storrie

## Abstract

**Background:** The contributions of platelet activation to thrombus formation during hemostatic bleeding cessation likely involve multiple activation states. However, the spatial and temporal distribution of platelets in these states has not been defined in a clot.

**Objectives:** To use single-platelet mapping of activation states within jugular vein puncture thrombi to determine how the spatial distribution of platelet state evolves during hemostasis.

**Methods:** Montaged, wide-area electron micrographs (EM) were taken at various time points, post-puncture, and annotated for platelet activation state. These classifications were mapped onto the images to identify regions of platelet activation and calculate neighbor associations. The importance of α-granule secretion was tested using VAMP8^-/-^ mice.

**Results:** mapping of platelet activation states at 1 min post-puncture showed extensive spatial intermixing of most platelet activation classes. No high-activation-state, platelet-rich core was observed in 5-min post-puncture thrombi, rather such platelets tended to be localized on the interior surfaces of thrombus vaults open to the circulation. Only at a later stage, 20 min post-puncture, was distinct clustering of high activation, degranulated, cytosol-rich platelets observed. These clusters localized to the central portion of the intravascular platelet-rich crown, and they were now inaccessible to the circulation. Counterintuitively, deletion of the primary platelet v-SNARE, VAMP8, increased the frequency and spatial clustering of highly activated, degranulated platelets in association with intra-thrombus vault surfaces at 5 min post puncture.

**Conclusions:** We conclude recent multi-activation state models can provide a realistic thrombus formation framework if linked together to encompass the dynamics of puncture wound formation.

## INTRODUCTION

Platelets are small anucleate cells that are about one-tenth as numerous as the most common cell type in blood, the red blood cell. The rapid recruitment of platelets to the exposed, collagen-rich adventitia of a puncture wound leads to the formation of a platelet-rich thrombus in a process that caps the wound from the extravascular side [1]. Hence, as indicated by our recent work [1,2], bleeding cessation comes not from simply plugging the hole, but from capping the outside and then filling the hole. Thrombus formation is accompanied by platelet activation. Morphologically, even within a bleeding, jugular vein puncture wound multiple platelet activation states are found [2]. These vary both in platelet shape, platelet-platelet adherence, and the presence or absence of granules. These changes follow rather than precede the recruitment to the puncture wound of discoid-shaped platelets resembling in morphology those in the circulation [2]. Considering the spatial separation of these thrombus-bound discoid platelets one from another, we suggested that they were tethered to one another by an elongated protein such as von Willebrand factor (VWF) in a process that we term “Capture and Activate” [2]. One prediction of a simple capture and activate hypothesis is the generation of a platelet aggregate in which platelets internal within the growing thrombus would be most activated and those at the thrombus periphery would be least activated. This is a prediction that, although different in reasoning, is like the classic “Core and Shell” hypothesis of thrombus formation proposed in the 2010s by others [3–6]. Neither hypothesis in its simplest form may fit the experimental data.

To test these hypotheses at the single platelet level, we determined platelet activation state distributions across a murine, jugular puncture wound model, at early and late time points. We also analyzed the effects of deleting the primary v-SNARE fusion protein involved in α-granule secretion [7] on resulting thrombus structure, bleeding cessation, and platelet activation state stratification. The ability to do this arises from the capacity of computerized, montaged, wide area transmission electron microscopy (WA-TEM) to yield high resolution, unbiased, 2D, platelet sampling across the full hundreds of microns width and height of a needle-induced, murine puncture wound [1,2]. Following manual annotation, a number representing the morphologically based activation state of each platelet profile can be assigned, the platelet X, and Y position of each platelet recorded, a multicolor map representing the distribution of each activation state, and a spreadsheet generated associating platelet state as a number with X, Y position of the respective platelets. From the spreadsheet summary data, for example, the abundance of individual activation states within the thrombus can be computed. Moreover, the spreadsheet supported the repeated sampling, on either a global scale or region of interest scale, of the frequency of platelet pairings across small distances within the thrombus. Platelets were grouped into activation classes based on their shape, granule status, and cytoplasmic content. These ranged from discoid to adherent platelets of various extents of granulation to granule-free platelets and finally to putative procoagulant platelets devoid of cytosol.

From this analysis, several significant outcomes emerge. 1) There is no highly activated core to the early thrombus; rather the emergence of any core of activated platelets is a late step in thrombus formation that has little spatial relationship to the exposed adventitia lining of the puncture wound hole. 2) Initially, there is very limited spatial segregation between different activation classes. 3) The bulk numerical distribution of platelets across activation class does not vary significantly over 20 minutes post puncture. 4) Region of interest analysis indicates the emergence of flow-dependent events in the distribution of discoid platelets, upstream versus downstream, in the periphery of the intravascular crown. 5) Significantly, a decrease in crown size at 20-min is associated with a shift in the distribution of high activation state platelets towards the center of the crown, an event that correlates well with thrombus compression and has the net effect of making these highly activated platelets inaccessible to circulating coagulation factors [see also, 1]. 6) VAMP8 granule fusion protein deletion had no obvious effect on bleeding cessation times, but it did affect thrombus structure leading to a thrombus that was leaky to red blood cells, less well formed, and perhaps paradoxically contained more highly activated platelets about peripheral thrombus surfaces, particularly those of internal cavities within the thrombus. In brief, any simple model including “Tether and Activate” requires extensions and modifications to account for the presumed signaling-induced changes in thrombus organization that occur over time. Presently, the complexity of puncture wound thrombus structure is best explained by a multiphase model that places various model elements into a timeline. Only such a model can explain the fact that the peripheral surfaces of vaults within a forming puncture wound thrombus are frequently lined by high activation platelets, rather than the low activation platelets expected by a simple Capture and Activate model.

## MATERIALS AND METHODS

### Mice and Reagents

All animal usage was approved by the relevant local Institutional Animal Care and Use Committees. Wild-type C57BL/6 or VAMP8^-/-^ male and female mice (8-12 weeks old) were used in equal numbers across the individual data sets. All reagents were of reagent grade and listed previously [1,2].

### Thrombus Preparation and Electron Microscopy

Jugular vein wounding was done with a 30 G needle (300 μm nominal diameter) and thrombi were fixed *in situ* at 1, 5, and 20 min, post injury, with 4% paraformaldehyde. For Wide -Area Transmission Electron Microscopy (WA-TEM), samples were processed for plastic embedding as previously described and stained with uranyl acetate and lead citrate post-embedding [1,2]. Automated montaged images were collected at 3.185 nm XY pixel size using SerialEM software (version 3.6, 32-bit) and visualized with 3DMOD software (version 4.9.13). Fine image blending was done with eTomo software (version 4.11.12). Image visualization was done with IMOD software (version 4,11). iMac Pro computers (MacOs 10.14, 5K display) were used, and images were displayed at various zoom factors ranging from 2% to 100%. The raw images were as large as 130,000 by 90,000 pixels. All software is from the Mastronarde group at https://www.colorado.edu/mcdb/resources/mastronarde-group.

For SBF-SEM, samples were stained with osmium tetroxide, uranyl acetate, and lead citrate pre-embedding [1 and references cited therein]. Segmentation and data analysis were performed as described [1]. The imaged thrombi were rendered in the following color scheme: vessel wall (blue), tightly adherent platelets (green), loosely adherent platelets (yellow), degranulated platelets (orange), and red blood cells (red).

### Datasets

The raw images of two of the 1 and 20-minute post-puncture thrombi (WA-TEM, 3.185 nm pixel size) were published previously at a low magnification [1,2]. These datasets are publicly available from the Electron Microscopy Public Image Archive (https://www.ebi.ac.uk/empiar/; EMPLAR ID accession code 10785, September 2021) and were used for the mapping analysis presented here. Additional datasets were generated to increase the robustness of our analysis. These will be deposited and publicly available as raw images. Upon acceptance of this manuscript. All mapping analysis, resulting quantitative analysis, and mapping imagery are unique and original to the present work.

### Platelet Activation State Mapping and Nearest Neighbor Analysis

Montaged WA-TEM images recorded in 24-bit, .mrc format were blended into a single image using eTomo (IMOD software package), binned (∼5x5) in 3DMOD, and converted to 8-bit grayscale images with FIJI software. Platelet activation states were annotated in the binned images. and displayed in 8-bit color using iVision-Mac software (32-bit software, BioVision Technologies, Inc., Exton, Pennsylvania). From the WA-TEM images, platelets were classified into five groups based on their morphology: discoid (blue); rounded, granulated (cyan); partially degranulation (green); degranulated cytosol-rich, but containing mitochondria (yellow); and devoid of internal contents (red). On the images, each platelet was scored, marked with a point in its center (centroid) which was given a unique identification number, and its X, Y coordinates were recorded. Platelet scoring was done manually by blinded scorers. The ‘Measure Segments’ command in iVision was used to evaluate the distributions of platelet states in the region surrounding each individual platelet. For each centroid, the state of all adjacent centroids, within a 2 µm radius, was tabulated. This was repeated iteratively for each platelet centroid in each field. The clustering of different platelet classes was calculated using the average numbers of each class associated with a given centroid and expressing that as a percentage of the total for that class. The percentages were graphed (y-axis) for each class of the centroid (1-5, x-axis) to denote how platelets in the different classes are grouped in a thrombus. Lower percentages, for example the discoid (blue) class, (see Figure 1 for example), indicate their even distribution with limited clustering. Higher values, for example the degranulated cytosol-rich, but containing mitochondria (yellow) class, indicate localized co-clustering of those types of platelets. Between 10,000 and 25,00 platelets were evaluated for each image set. Calculations were done with Excel, see Supplemental Materials for example.

**Figure 1.**
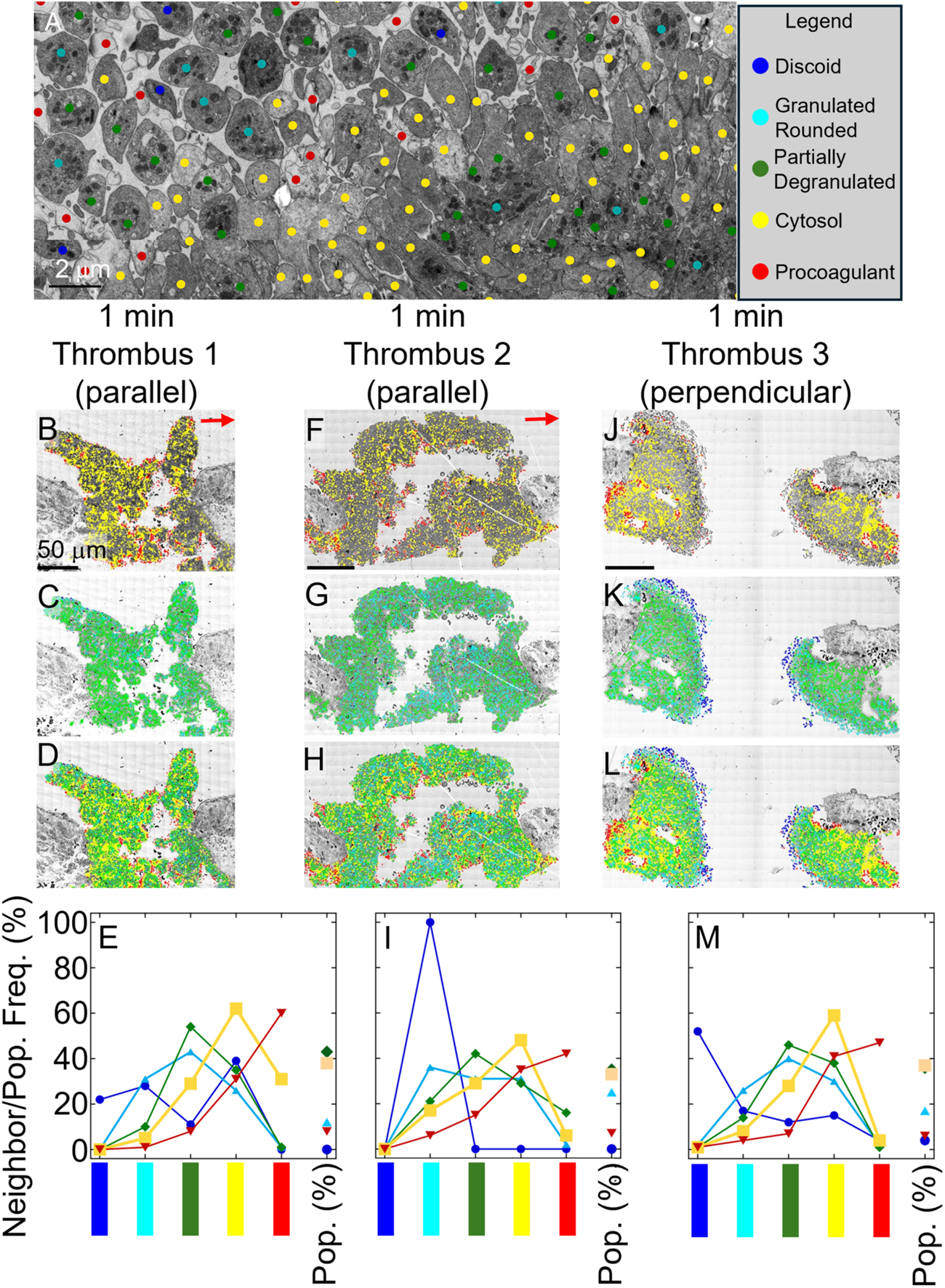
Distribution of Platelets in Venous Puncture Thrombi, 1 min post injury. Individual platelets were scored based on morphology in the WA-TEM images of thrombi formed in the first minute after a puncture injury of a mouse jugular vein. **A**, Examples of the 5 distinct platelet states used for scoring. Color coding is described in the Methods. **B-I,** Scored images and quantification (E and I) for thrombi #1 and 2 respectively, which is sectioned mid-thrombus, on an axis parallel to flow. **J-M,** Scored images and quantification (M) for thrombi #3, which is sectioned mid thrombus on an axis orthogonal to flow. Neighbor relationships within a 2 μm radius from the individual center point; far right column, population % of each class. Red arrows mark the direction of blood flow. Scale Bar = 50 μm.

## RESULTS

### Ultrastructural Analysis of Puncture Wound Thrombi

Previously, we used serial block face-scanning electron microscopy (SBF-SEM) supplemented by limited WA-TEM to render full thrombus volumes of the jugular vein, puncture wound thrombi produced by a 30 G needle, 300 µm nominal diameter [1,2]. Samples were fixed *in situ* at 1, 5, and 20 min post puncture. These experiments provided evidence that highly activated platelets in 1 and 5 min samples were located in a thin layer lining platelet-free cavities within the forming thrombus. By 20 min post puncture, cavities had shrunk, and the overall thrombus volume decreased relative to that at 5 min. Because of the relatively low resolution of these SBF-SEM-based experiments, 100 nm typical pixel size, we gained little knowledge of the activity state of individual platelets within the forming thrombi and hence could make at most only general inferences about platelet morphology, putative platelet activation class distribution, and how signaling might change locally within the forming thrombi.

To overcome these limitations and still be able to place sections precisely in 3D space within the thrombi, we attempted to score platelet activation levels from SBF-SEM images taken at 20 nm pixel size, every 20 μm across the puncture wound. Unfortunately, the SBF-SEM images after required pre-analysis binning lacked the detailed resolution needed to score platelet activation at the single cell level, to map platelet stratification, and to determine platelet-platelet neighbor to neighbor relationships across full thrombus planes. As an alternative, we turned to computer-assisted, manual platelet activation state annotation using montaged WA-TEMs taken at 3.185 nm X,Y pixel size. The original montaged images were converted to 8-bit grayscale images and scaled from the original 20 to 30-gigabyte size to approximately 1 gigabyte or less, ∼5x5 binning, because of 32-bit limitations built into the annotation software used. Images were displayed on a 5K screen and magnified for scoring. As shown in **Figure 1A**, this was sufficient to support platelet classification into five activation classes: discoid (blue); rounded, granulated (cyan); partially degranulation (green); degranulated cytosol-rich, but containing mitochondria (yellow); and procoagulant, devoid of internal contents (red). Labeling and color choice was made to indicate progressive activation classes. Between 10,000 to 25,000 platelet profiles were scored for each WA-TEM montaged cross-section. The initial platelet “click” was manual, i.e., to the eye of result-blinded scorers. That click automatically linked activation classification and XY position for each platelet to a map and a spreadsheet.

### Initial post puncture wound thrombi show little platelet activation state stratification

Three individual puncture wounds were scored,1 min following jugular vein puncture. In none of the three examples was the puncture wound completely closed. The TEM images were acquired approximately 50% into the puncture wound. Sectioning was done manually and hence placement within the thrombus was somewhat arbitrary. Two of the three samples were sectioned parallel to flow (left to right in images) and one sample was sectioned perpendicular to flow. The individual maps shown in Figure 1 are, as in later examples, presented in 3-panel variations: hot colors (red and yellow, high activation), cold colors (blue, cyan, and green, weak to moderate activation), and as full 5-color images. The highest activation class, procoagulant, devoid of cytosol (red) was located adjacent to exposed adventitia or on exposed peripheral surfaces, particularly those of internal cavities within the forming thrombus (see **Figure 1B, F, J**, S**upplemental Figure 1**, 5 colors, full page image). Few procoagulant platelets were found internally within the platelet-rich aggregates. In contrast, cytosol-rich, degranulated platelet profiles (yellow) were found over much of the forming thrombi with the exceptions of some internal areas (**Figure 1B, F)** that were devoid of cytosol-rich, degranulated platelets. Discoid platelets (blue) were especially numerous on the whole exposed surfaces of the third cross-section cut perpendicular to flow (**Figure 1K**). As shown in **Figures 1C, G, and K**, moderate activation platelets, especially those with varying degrees of granulation (cyan and green) were found over most of the 1 min thrombi. In total about 80% of the platelets scored fell into these two classes of granule-positive, rounded, adherent platelets. The higher activation of the peripheral areas of the 1 min thrombi was consistent with previous qualitative results [1]. The fact that there was a little gradation in platelet state within the thrombus aggregates suggests that conversion from a discoid-shaped platelet to a granule-positive, more rounded state occurs quickly.

To give quantitative comparisons of the global, nearest neighbor activation state, we used XY spatial matching to determine neighboring platelet classification within a 2 μm radius (centroid to centroid) of any given platelet (**Figure 1E, I, M**). On a random basis, we expected that neighbor pairing would be directly proportional to the frequency of any classification state. For example, platelets showing some degranulating, adherent, rounded morphology (green) should pair randomly at a frequency of 40-45% with each other. Similar pairing frequencies would be expected for rounded, granulated, adherent platelets. Quantitatively, each class self-associated with a frequency ∼10 percentage points higher than expected. In contrast, the two lowest activation classes, discoid (blue) and rounded, fully granulated (cyan), showed a stronger preference for patterned association, often two-fold or more with, in one case, nearly 100% of discoid platelets being associated with rounded, fully granulated platelets. Strikingly, procoagulant-like platelets showed a strong self-association preference, 5-fold or higher.

Our 1-min post puncture mapping results point to a mixed salt and pepper situation in major activation classes distribution. There was a limited tendency of major activation classes to self-associate beyond that expected by chance. Visually, significant local examples of self-association were those of discoid platelets within the puncture hole of a perpendicular section or more broadly that of rounded, fully granulated platelets, and procoagulant platelets proximal to the exposed adventitia. We suggest that these results support a “Capture and Activate” model of initial platelet recruitment in which initial adherence at the collagen-rich adventitia produces a procoagulant state. Repeated platelet recruitment within the puncture hole leads to subsequent activation that does not progress past partial degranulation as indicated by the morphologically identifiable large granule class, α-granules. In net, these results present little, if any, evidence for local patches or larger areas of highly activated platelets within early, time-staged puncture wound thrombi that might approximate a Core as defined in the “Core and Shell” model [3–6]. Note: No effort was made to quantify dense granules within the 1-min or later thrombus cross sections. The incidence of dense granules within platelet cross-sections is too small to make their quantification meaningful [8].

### Mature post-puncture thrombi show centralized concentrations of highly activated platelets and flow-dependent stratification within the intravascular region

To test whether these outcomes hold across the late-stage puncture wound thrombi (previous results indicated a considerable remodeling and compression [1], we applied the same analytical approach to WA-TEM montaged cross sections of 20-min post puncture thrombi. Again, three examples, two parallel and one perpendicular to flow, were analyzed. As apparent from even visual inspection, there appears to be a significant concentration of cytosol-rich, degranulated platelets (yellow color) within the central, internal portion of each of the three individual thrombi. The first parallel example and the perpendicular example also showed concentrations of degranulated platelets in exposed adventitial collagen-rich regions (**Figure 2A, E, I, C, G, K**). Furthermore, when the map display was restricted to the three low activation classes (blue, cyan, and green colors), a distinct flow-dependent difference in low activation state mapping became even more apparent. The downstream portion of the crown in both parallel cut examples qualitatively showed a higher number of discoid (blue color) and rounded, high granulation platelets (cyan color) to the downstream side of the intravascular crown versus the upstream side (**Figure 2B, F**). As expected, such a difference was not apparent in the perpendicular cut cross-section example, **Figure 2J**. In all three examples, irrespective of flow, the more peripheral portions of the intravascular crown appeared relatively deficient in partially degranulated platelets (green color).

**Figure 2.**
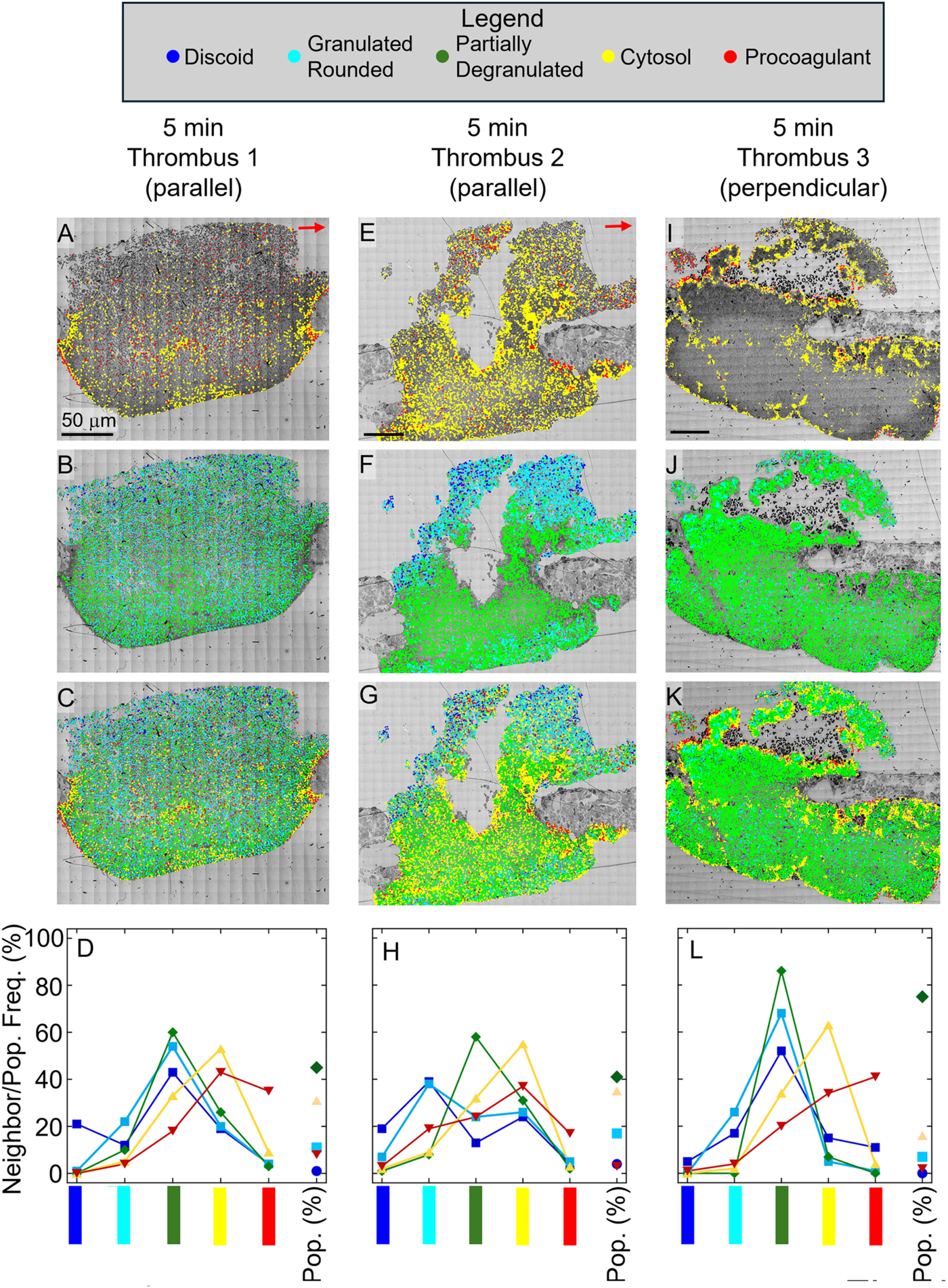
Distribution of Platelets in Venous Puncture Thrombi, 5 min post injury. Individual platelets were scored based on morphology in the WA-TEM images of thrombi formed at five minutes after a puncture injury of a mouse jugular vein, when bleeding had just ceased. **A-H,** Scored images and quantification (D and H) for thrombi #1 and 2 respectively, which is sectioned mid-thrombus, on an axis parallel to flow. **I-L,** Scored images and quantification (L) for thrombi #3, which is sectioned mid thrombus on an axis orthogonal to flow. Neighbor relationships within a 2 μm radius from the individual center point; far right column, population % of each class. Red arrows mark the direction of blood flow. Scale Bar = 50 μm.

To quantify these outcomes, we took two approaches. The first was, as before, global quantification of neighbor pairings over the cross sections (**Figure 2D, H, L**). These showed little difference with those of the 1-min post puncture thrombi (**Figure 1E, I, M**) with the most activated class, procoagulant platelets (red color), which failed to show as much self-association suggestive of clustering. As quantified in **Figures 1 and 2**, there is no significant difference in global neighbor pairings between 1-min and 20-min post puncture. Likewise, as shown in **Figure 3**, there are no significant differences globally between the 1 min and 20 min post puncture, mid thrombus cross sections in total number of platelets and individual activation class percents. We conclude that the qualitative upstream-downstream differences noted above are obscure when the global values are considered. To highlight the local upstream-downstream differences apparent in the maps qualitatively, we used region of interest (ROI) analysis. In **Figure 3D**, ROI quantifications are shown for the respective areas superimposed on the 5-color maps in **Figure 2**. The platelet images within the respective ROIs are shown in **Supplemental Figure 2**. These quantifications validate the qualitative outcomes presented in the preceding paragraph.

**Figure 3.**
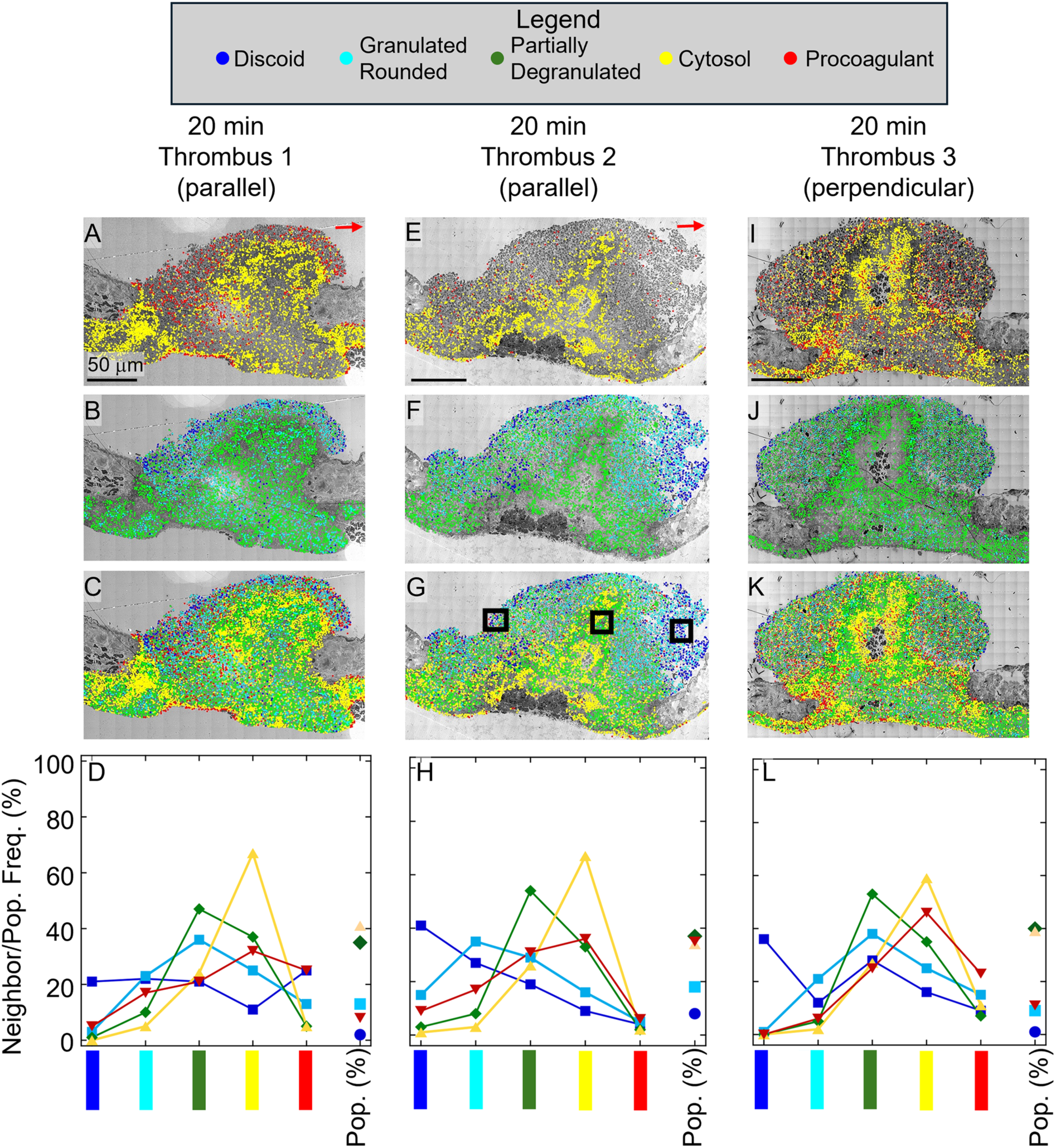
Distribution of Platelets in a Venous Puncture Thrombi, 20 min post injury. Individual platelets were scored based on morphology in the WA-TEM images of thrombi formed at 20 minutes after a puncture injury of a mouse jugular vein. **A-H,** Scored images and quantification (D and H) for thrombi #1 and 2 respectively, which is sectioned mid-thrombus, on an axis parallel to flow. **I-L,** Scored images and quantification (L) for thrombi #3, which is sectioned mid thrombus on an axis orthogonal to flow. Neighbor relationships within a 2 μm radius from the individual center point; far right column, population % of each class. Red arrows mark the direction of blood flow. Scale Bar = 50 μm. The black boxes in G mark the three regions of interest that were analyzed further, see Figure 4.

### Deleting the primary v-SNARE for platelet granule fusion does not affect bleeding but does affect thrombus architecture

To address the effect of perturbing α-granule secretion on thrombus formation, we compared thrombus structure and mapping outcomes between Control and VAMP8 knockout mice at 5 min post puncture. We chose to perturb α-granule release kinetics by using VAMP8 knockout mice as it is the primary protein implicated in α−granule fusion and its deletion slows and depresses α-granule release [7,8]. By 5 min post puncture, the thrombus is expected to show an intravascular crown with typically extensive vaults and substantial extravascular platelet accumulation that caps the puncture hole and prevents further bleeding [1,2]. The images from wild-type control mice shown in **Figure 4** are consistent with those expectations. A substantial extravascular cap was present in each of the four examples (**Figure 4** and **Supplemental Figure 3**). Rounded platelets displaying some degranulation (green color) were common, almost 80% of total platelets in one example (**Figure 4J, L**). Higher activation state platelets (e.g., cytosol rich, degranulated, yellow color) were generally distributed across the thrombi with some concentration at the periphery of internal cavities. These results contrast with those shown in **Figures 5 and 6** for 5 min post-puncture VAMP8 knockout mice thrombi.

**Figure 4.**
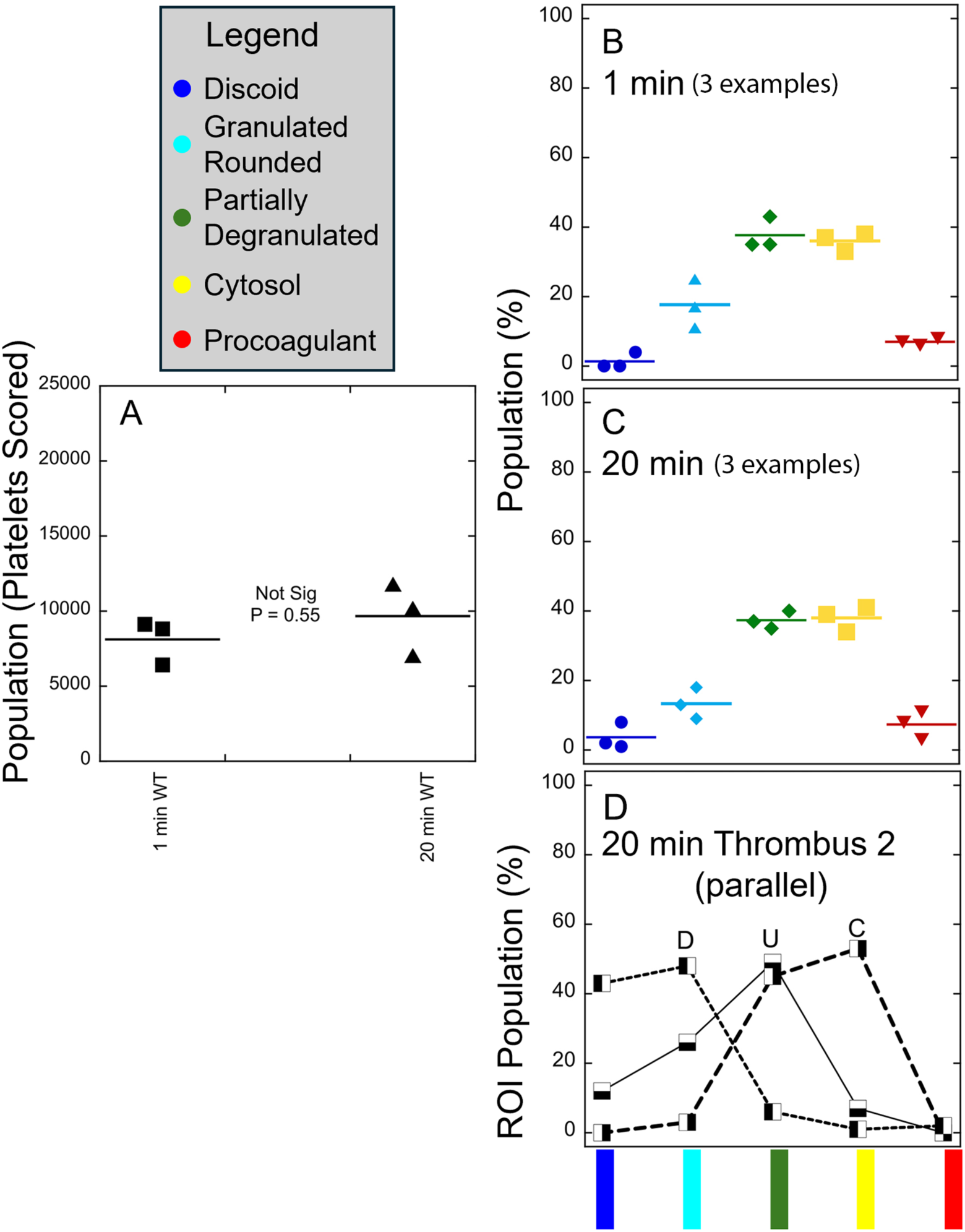
Quantitation of thrombus size (A), global activation class size (B,C), and intravascular region of interest analysis (D,upstream, central, and downstream, black squares in Figure 3G) reveal similar overall thrombus size and percent occurrence of each activation at 1 versus 20 min despite the increased spatial concentration of activation classes. A, Population number of platelets scored in 1 and 20 min thrombi. B, C, Global activation state distribution between activation classes, 1 and 20 min post-puncture. D, Region of interest illustrative outcome for intravascular regions within a 20 min thrombus.

**Figure 5.**
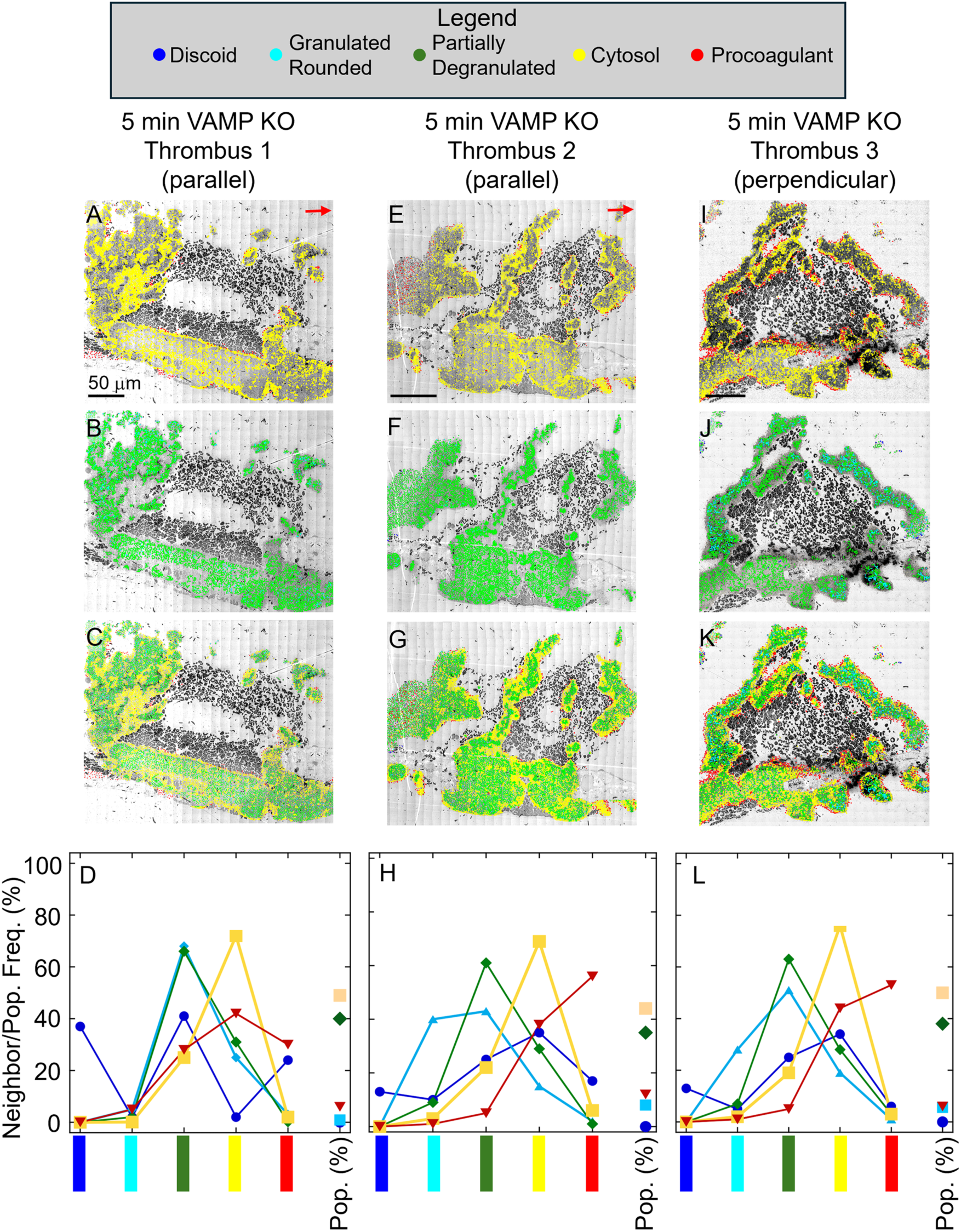
Distribution of Platelets in a Venous Puncture Thrombi, 5 min post injury when secretion is reduced by deletion of VAMP-8. Individual platelets were scored based on morphology in the WA-TEM images of thrombi formed at 5 minutes after a puncture injury of a VAMP8^-/-^ mouse jugular vein. **A-H,** Scored images and quantification (D and H) for thrombi #1 and 2 respectively, which is sectioned mid-thrombus, on an axis parallel to flow. **I-L,** Scored images and quantification (L) for thrombi #3, which is sectioned mid thrombus on an axis orthogonal to flow. Neighbor relationships within a 2 μm radius from the individual center point; far right column, population % of each class. Red arrows mark the direction of blood flow. Scale Bar = 50 μm.

**Figure 6.**
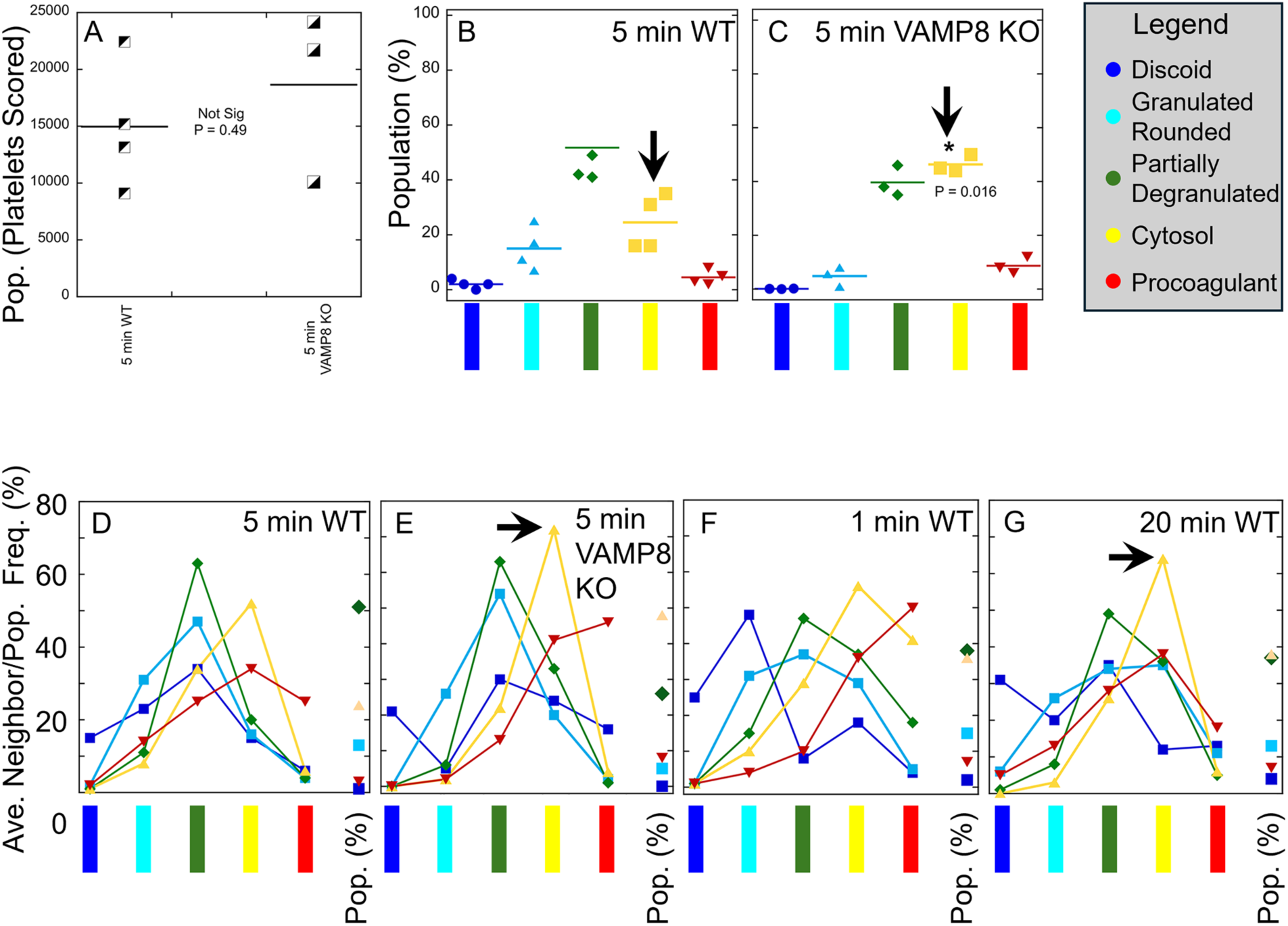
Quantitative Comparisons of Platelet State Distribution in Wild-type and VAMP-8^-/-^ mice. Pairwise quantification of platelet number and classes in wild type versus VAMP8-/-mice and between 1 and 20 min post-puncture thrombi in Wild-type mice. **A-C,** Population number and percent and various platelet activation classes in 5 min post-puncture jugular vein wounds. **A,** Total platelet count in wild type and VAMP8 knockout thrombi when mid-thrombus are not significantly different. **B, C**, The population percent of degranulated cytosol-containing platelets, high activation state, increases significantly in VAMP8 knockout thrombi (arrowheads). **D, E**, Average neighbor pairings in 5 min post-puncture wild type versus VAMP8 knockout thrombi trend towards a tighter pairing of degranulated cytosol containing platelets (yellow) one with another in the VAMP8 knockout thrombi (**E**, arrowhead). Similarly, the pairing trends are higher in wildtype 20 min versus 5 min post-puncture thrombi (**G**, arrowhead).

As anticipated deletion of VAMP8 [7,8] had minimal to no effect on puncture wound bleeding time in the jugular model (**Figure 5A**). However, by SBF-SEM across the full width of the wound hole, the thickness of the extravascular cap was thin in the KO (**Figure 5B**, 20 nm pixel raw image every 20 μm). Moreover, rendering of the images taken every 200 nm at 100 nm pixel size revealed small cracks in the extravascular cap when the obscuring RBC layer in the surface rendered image was deleted (**Figure 5C** versus **5D,** circled area). These small cracks could lead to RBC leakage through the extravascular cap of the relatively disorganized KO thrombi (**Figure 5E**, full intravascular surface rendering). By WA-TEM (2% Zoom, **Figure 5F**), the extravascular cap was indeed found to be leaky to RBCs and intravascularly more disconnected than expected for a 5-minute thrombus.

All three WA-TEM examples mapped in **Figure 6** showed RBC leakage and appeared disorganized relative to the wild-type examples shown in **Figure 5**. Visually each exhibited a cavity-filled interior filled with RBCs that was open to the intravascular vessel lumen. Surprisingly, each thrombus also was marked by a high level of degranulated platelets (yellow color, **Figure 6A, E, I, C, G, K**) lining peripheral surfaces and enclosing less activated interior platelets. Quantitatively, the frequency of degranulated platelets almost doubled relative to that of 5 min wild-type thrombi (**Figure 7**). The increase in degranulated platelet frequency was accompanied by a decrease in the frequency of moderately activated, partially degranulated platelets abundant in the thrombus interiors. Interestingly, VAMP8 KO had almost no effect on the number of platelets per cross-section compared to control, wild type 5 min thrombi (**Figure 7**). Both wild type and VAMP8 KO 5 min thrombi exhibited a roughly two-fold increase in platelet number per cross-section when compared to either 1-min or 20-min post-puncture thrombi (**Figure 3** versus **Figure 7**). In conclusion, we speculate that the high level of platelet degranulation observed in the VAMP8 KO thrombus is due to the relatively chaotic state of thrombus organization in the KO and supports an accumulation of vascular stimulatory factors within the extensive cavities of the KO thrombi. Alternatively, the accumulated RBCs could be a factor [9–11].

**Figure 7.**
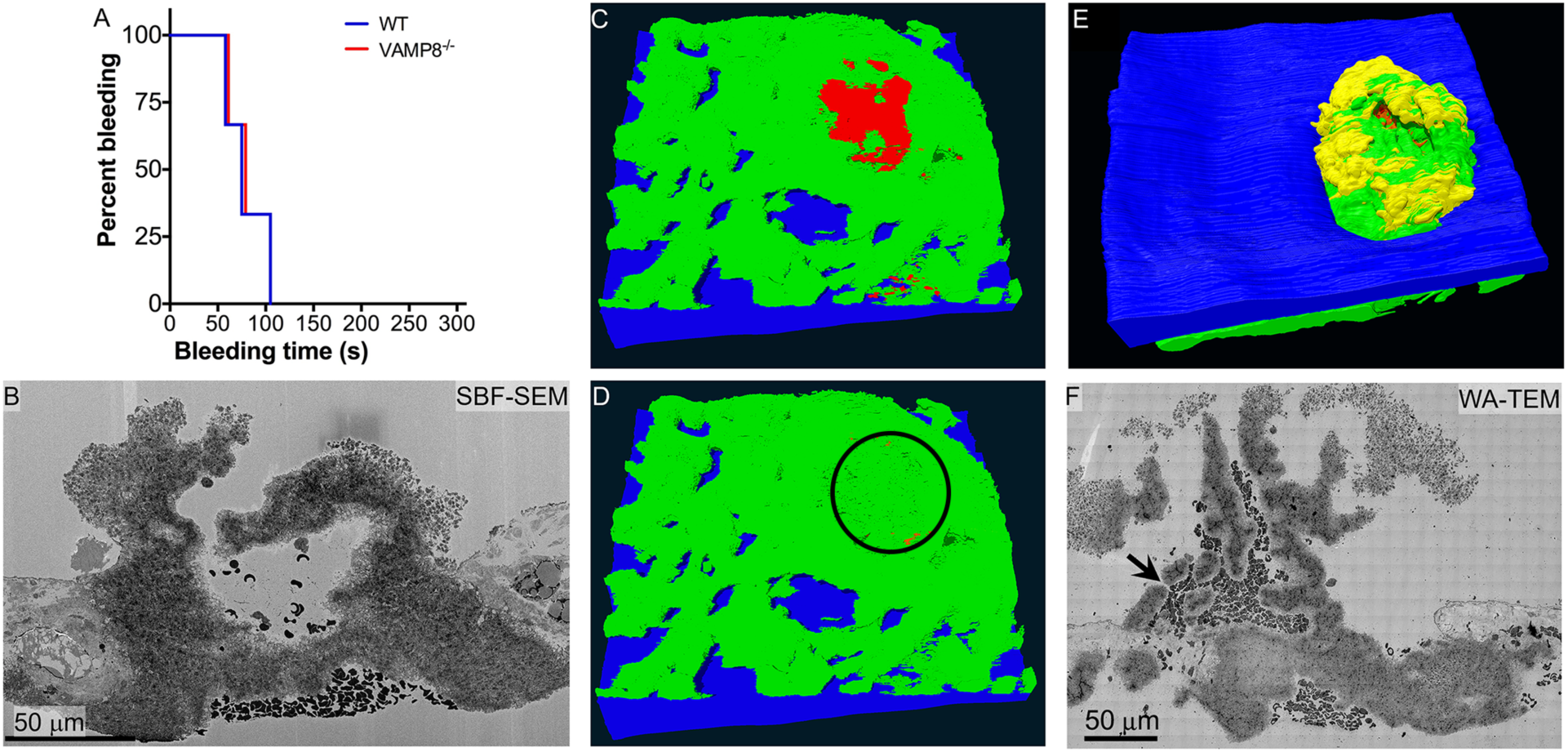
Venous Puncture Thrombi Structure, 5 min post injury when secretion is reduced by deletion of VAMP-8. **A**, Bleeding cessation times for wild type and VAMP8^-/-^ jugular vein puncture wounded mice. **B-E**, Raw and rendered images of a 5-min, VAMP8 knockout puncture wound thrombus. In C-E, the color coding is RBC, red; tightly adherent platelets, green; and loosely adherent platelets, yellow. C, D, a rendered view of the extravascular surface. Circle in D encloses a set of small openings in the extravascular side of the thrombus that could be leakage points for the extravascular accumulation of RBCs. E, view of the intravascular surface of the rendered thrombus. F, WA-TEM (3.185 nm raw pixel size, top intravascular; bottom extravascular) reveals small openings in the knockout thrombus through which RBCs appear to leak (e.g., black arrow). The light, peripheral areas in the raw image in F are shown in mapping to be highly activated platelets that degranulate and continue to contain cytosol components. Scale Bar = 50 μm.

## DISCUSSION

The process of platelet aggregation is fundamental to thrombus formation, be it hemostatic or thrombotic, and requires at least some degree of platelet activation. Circulating platelets have little to no tendency to aggregate. We scored *in vivo* individual platelet activation states and aggregation to test this premise in a model of thrombus formation, i.e., a mouse puncture wound paradigm of profuse bleeding. We used two complementary volume EM techniques, WA-TEM and SBF-SEM, to visualize platelet activation and computer-assisted methodology to convert annotated ultrastructure into maps of individual platelet activation states within forming jugular vein puncture wound thrombi and then performed quantitative analysis of neighboring state pairing within the thrombi. In brief, we generated the experimental outcomes needed to test the proposed roles of platelet activation and α-granule secretion in current models of thrombus formation. For example, the “Core and Shell” model proposes a distinct, multi-platelet deep layer of highly activated platelets at the thrombus/exposed adventitia interface that is postulated to anchor any further thrombus formation [3–6]; such is an outcome based chiefly on fluorescence light microscopy and scanning electron microscopy [e.g.,5], techniques which tend to detect at best only platelets at the thrombus periphery and fail to reveal the state of individual platelets. On the other hand, the “Cap and Build”/” Capture and Activate” models propose that thrombus formation is initiated at small, nucleation sites along the exposed adventitia and grows by a tethered capture of circulating platelets which then activate progressively to varied states within the forming thrombus [1,2]. That set of projections is based chiefly on volume EM techniques in which individual platelets are visualized at comparatively low resolution, typically 20 to 100 nm pixel size at the electron microscope, across full puncture wound diameters.

We conclude that WA-TEM, an example of volume EM approaches, is sufficient to lead to an accurate thrombus mapping across tens of thousands of platelet profiles within an individual thrombus when paired with manual annotation of platelet activation state followed by a computer-assisted map display and computation of platelet activation state neighborhoods. Most of our analysis started from montaged 2D WA-TEMs across full mid-thrombus cross sections at a typical minimum raw pixel size of 3.185 nm and in some cases even smaller. That meant that after binning to accommodate 32-bit image analysis software we were working with images containing pixels of 12-15 nm size. Hence, the internal features of the thrombus and platelets were revealed in detail. Our analysis revealed consistent results within individual time points and consistent differences across sequential time points. It was sufficient to reveal a significant mutational disruption of thrombus structure and platelet activation due to knockout of the primary v-SNARE fusion protein, VAMP8, implicated in α-granule release [7] even though bleeding times indicated no significant effects. Furthermore, the platelet activation classes observed in these primarily 2D experiments are entirely consistent with those found in our ongoing 3D region of interest assessment of morphological variation in platelet activation state, albeit, at an analysis level of 100 platelets in 3D versus hundreds of thousands of platelets as in the current work (Madhavi Ariyathne et al., manuscript in preparation).

These studies lead to three major outcomes. First and perhaps surprising, most platelets, 50-70%, in a forming thrombus, be it one scored at 1, 5, or 20 min post puncture, were granulated, i.e., contained α-granules, the protein containing, major secretory organelle of platelets and the most numerous and readily detected granule within platelets. These granulated platelets were in the interior of the platelet aggregates presumably as the result of a tether and activation process in which such interiorly located platelets are exposed to limited signaling levels. Second and consistently, there was no highly activated platelet core present early in a 1 min post-puncture thrombus. Any grouping of platelets to form what might be termed a highly activated core occurs late, several min after bleeding cessation, with most of the grouping distal from the exposed adventitia/wound thrombus interface. The clustering of highly activated platelets was located primarily within the central interior of a 20-min post-puncture thrombus and was likely due primarily to clot compression, presumably due to fibrin contraction [see also 1]. Third, over time, 1, 5, and 20 min, the spatial distribution of platelet activation states evolves. Although the relative distribution of platelets across activation classes stays constant, the spatial pattern changes, Initially, at 1 min post-puncture, there is little spatial separation between classes whereas at 5 min post-puncture there is a concentration of apparent, high activation state degranulated platelets along circulation exposed surfaces of vaulted cavities within the thrombi, Later, at 20 min post-puncture, as the vaults appear to be compressed, a central grouping of highly activated platelets becomes evident. In brief, there is thrombus remodeling/spatial evolution at the level of the grouping of individual platelet activation states. Functionally, this evolution may well be quite important, intermixed with the high activation, degranulated platelets, we found ballooned, apparent cytosol free platelets. Morphologically, these platelets have the properties of procoagulant platelets. Their exposure to the circulation at 5 min may well contribute significantly to the activation of coagulation factors. If so, the later vault compression and the placement of highly activated platelets within the interior of the thrombus could well be a thitherto unrealized mechanism for a thrombus structure that self-limits further coagulation factor activation. This is an interesting but speculative hypothesis for now that is deserving of further testing.

Interestingly, the frequency of highly activated platelets was higher in 5 min post puncture thrombi from VAMP8^-/-^ mice. These platelets were concentrated on the exposed vault surfaces of VAMP8 knockout thrombi. VAMP8, the primary v-SNARE, for α-granule release, has been shown to produce a phenotype in which α-granule secretion is delayed [7,8]. We propose that disrupted thrombus structure in the knockout results in greater exposure of vault peripheral platelets to a higher soluble signal intensity and hence greater activation. Yet, within the interior of the thrombus aggregates, the level of signal intensity is too low to result in general platelet degranulation. Indeed, even under these conditions, most platelets are not degranulated. In conclusion, we suggest a functional explanation for the behavior of platelets in these examples of puncture wound thrombus formation, namely as putative, self-limiting in exposure, substrates for coagulation factor action and ask whether these phenomena occur in arteries as well as in vein. We propose a multi-phase model that separates thrombus formation into sequential, but overlapping, phases and consolidates our previous “Cap and Build” and “Capture and Activate” models [1,2] into what might be termed a “Tether and Build” model in which consolidation of thrombus volume limits the exposure of degranulated palates to the circulation (**Figure 8**). Initially, tethered capture of discoid platelets from the circulation leads to the progressive activation of the captured platelets within-/-the forming thrombus (1 min post-puncture). That places these platelets within the interior of the thrombus, a situation that limits soluble signal penetration and leads to at most partially degranulated platelets. With time, a Build phase produces vaulted chambers inside the thrombus, and within the vaulted chambers soluble signals accumulate. The accumulation of these postulated soluble signals produces degranulation of peripheral platelets lining the vaults (5 min post-puncture). These surfaces are exposed to intravascular circulation and potentially provides membrane substrata for rapid activation of coagulation factors. That is later followed by thrombus compression and the compression of vaults to give a thrombus interior relatively rich in highly activated platelets (20 min post-puncture). These putative procoagulation platelets are now no longer accessible to the circulation. Overall, this model is based on structural data from the jugular vein. However, our one published arterial example, a femoral artery puncture wound thrombus [1, see Supplemental Figures at journal website], does present an outcome structural consistent with our jugular vein-based model.

**Figure 8.**
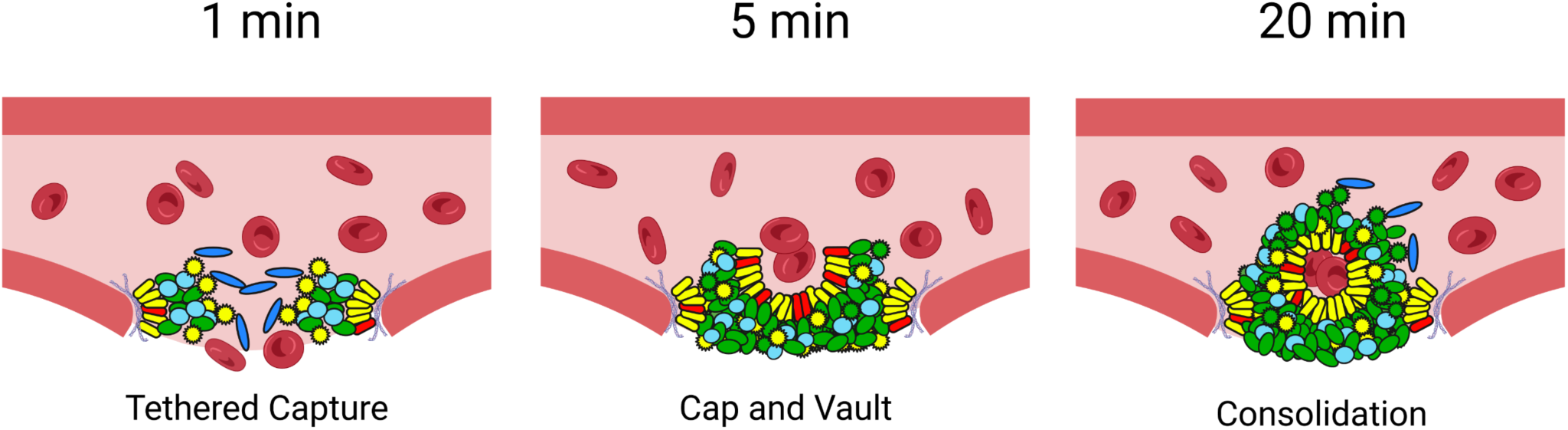
Multiphasic model for jugular vein puncture wound formation. The model is based on the results presented here and in two previous publications [1.2]. The platelet color coding system is consistent with the classification used in Figures 1-6.

## Supporting information

Supplemental Analysis Guide

Supplemental Template Example

## Role of Authors

Sung Rhee and Irina Pokrovskaya contributed equally to experimental implementation with major roles in data gathering. Kelly Ball was responsible for puncture wounds and all animal surgeries at UAMS. Jeffrey Kamykowski supported electron microscope sample preparation and electron microscopy at UAMS. All SBF-SEM was done by the Leapman laboratory at NIBIB with Oliver Zhao responsible for experimental implementation/data analysis steps and Maria A. Aronova and Richard Leapman being major contributors to experimental design and supervision. Irina Pokrovskaya led all electron microscope sample preparation and WA-TEM. Irina Pokrovskaya was responsible for data validation and coordination between the Storrie and Leapman laboratories. Richard Leapman, Sung Rhee, and Brian Storrie had major responsibilities for experimental design, data quality control, and manuscript preparation. VAMP8 knockout mice originated in the laboratory of Dr. Sidney W. Whiteheart at the University of Kentucky.

## Essentials

Though hemostasis biochemistry is well defined, the spatial/temporal distribution of individual, activated platelets in a thrombus is less characterized.

Detailed, wide-area electron microscopy and quantitative morphometrics map heterogeneity of platelet activation states in venous thrombi.

Initially, activated platelets are distributed uniformly and are folded into the interior as the thrombus matures.

Defective platelet secretion affects the consolidation of mature thrombi.

## Acknowledgments

Thanks are given for what was truly a team effort over several years. Gift puncture wound samples from Dr. Tim Stalker at the University of Pennsylvania in association with Dr. Lawrence F. Brass were significant contributions to initiating this work. Work at UAMS was supported by HL119393, HL119393, and GM155519 to BS. Work at the University of Pennsylvania was supported by HL040387 and HL120846 to TJS and LFB, respectively. Work at the University of Kentucky was supported by HL150818 to SWW. The Leapman laboratory was supported by the intramural program of NIBIB at the National Institutes of Health, Bethesda, MD.

## SUPPLEMENT FIGURES

An example spreadsheet analysis is available upon request as a separate file.

**Supplemental Figure 1.**
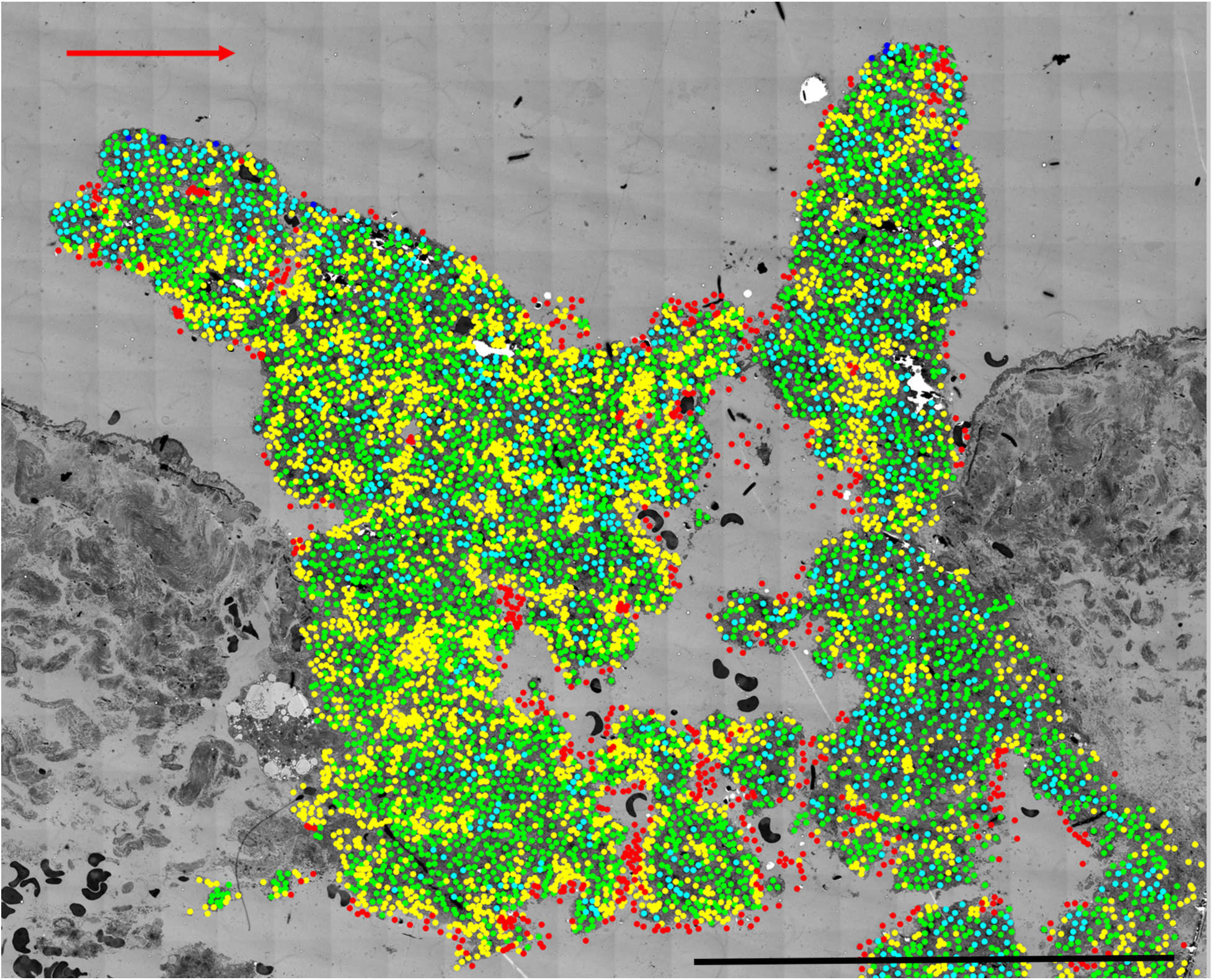
An enlarged image of 1 min 5 color map, Example 1. Flow left to right, red arrow. Bar = 100 μm

**Supplemental Figure 2.**
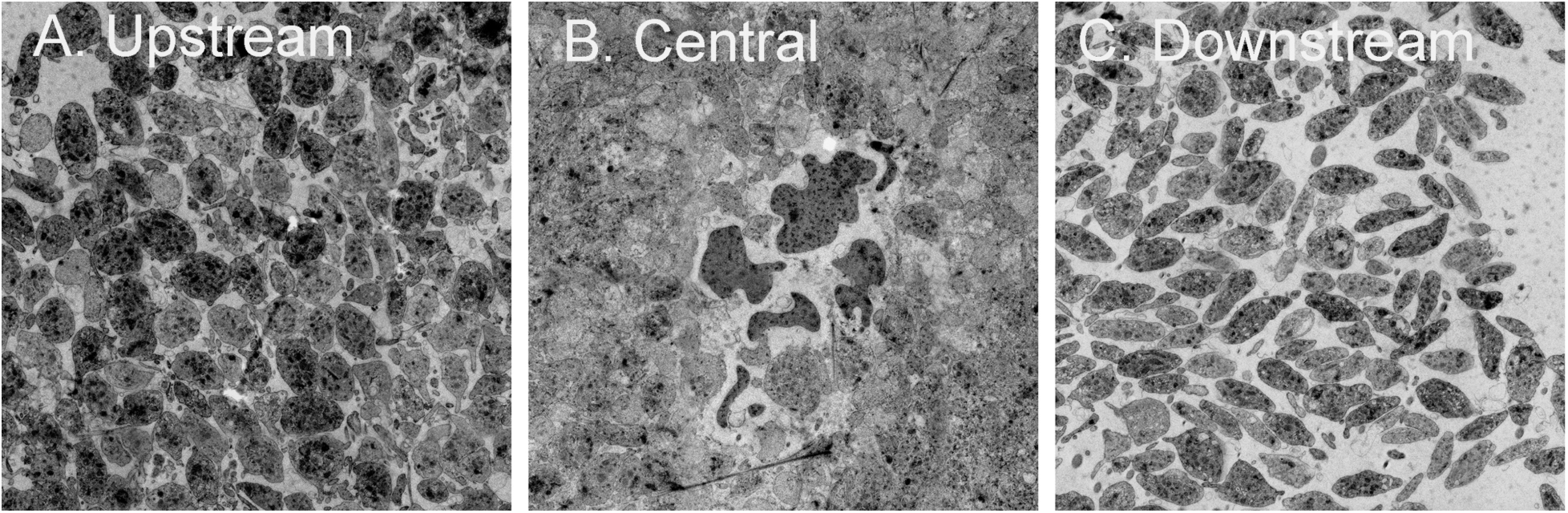
Raw image of platelets within Upstream, Central, and Downstream region of interest samplings of the intravascular crown of a 20 min post-puncture jugular vein thrombus.

**Supplemental Figure 3.**
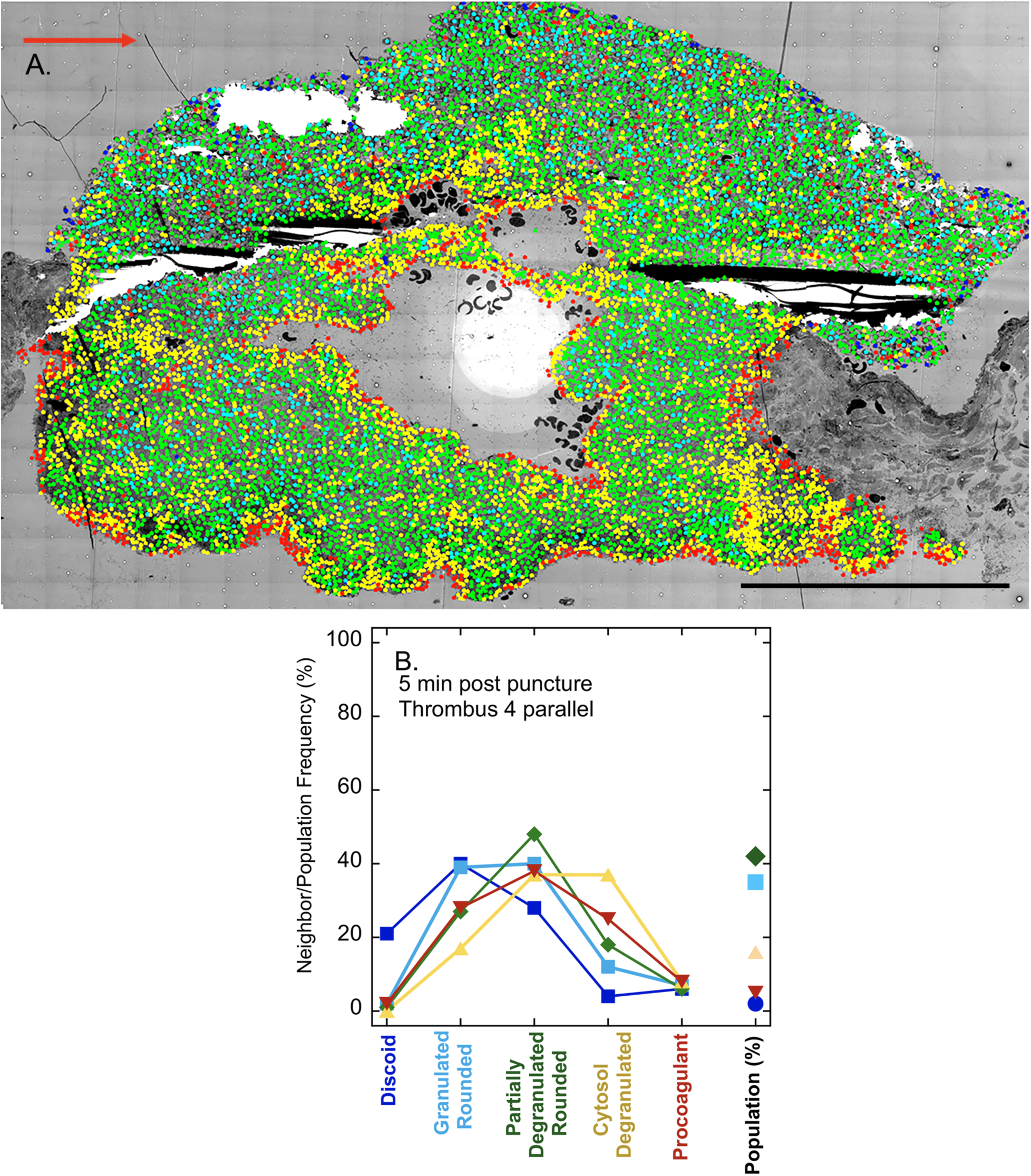
Five-color map and neighbor pairings for 5 min post-puncture thrombus. Note that there is some cracking during the preparation and missing frames in the pre-blended image set. Scale bar = 100 μm.

## REFERENCES

1. Rhee SW, Pokrovskaya ID, Ball KK, et al. Venous puncture wound hemostasis results in a vaulted thrombus structured by locally nucleated platelet aggregates. Commun Biol 2021;4:090. doi: 10.1038/s42003-021-02615-y.

2. Pokrovskaya, et al. Tethered platelet capture provides a mechanism for restricting circulating platelet activation to the wound site. Res Pract Thromb Haemost. 2023 Jan 23;7(2):100058. doi: 10.1016/j.rpth.2023.100058. eCollection 2023 Feb.

3. Stalker TJ, Traxler EA, Wu J, et al. Hierarchical organiation in the hemostatic response and its relationaship to the platelet signaling network. Blood 2013;121:1875–1885.

4. Stalker TJ, Welsh JD, Brass LF. Shaping the platelet response to vascular injury. Cur Opin Hematol. 2014;21:410–417.

5. Tomaiuolo M, Matzko CN, Posentud-Fuentes I, Weisel JW, Brass LF, Stalker TJ. Interrelationships between structure and function during the hemostatic response to injury. Proc Natl Acad Sci USA. 2019;116:2243–2252.

6. Brass LF, Diamond SL, Stalker TJ. Platelets and hemostasis: a new perspective on an old subject. Blood Adv. 2018;1:5–9.

7. Ren Q, Kalani Barber H Crawford G, Karim ZA, Zhao C, Choi W, Wang C-C, Hong W, Whiteheart SW. Endobrevin/VAMP-8 is the primary v-SNARE for the platelet release reaction. Mol Cell Biol. 2007 Jan;18(1):24–33. doi: 10.1091/mbc.e06-09-0785. Epub 2006 Oct 25.

8. Pokrovskaya I, Joshi S, Tobin M, et al. SNARE-dependent membrane fusion initiates α-granule matrix decondensation in mouse platelets. Blood Adv. 2018;2:2947–2958..

9. Barr JD, Chauhan AK, Schaeffer GV, Hanssen JS, Motto DG. Red bloot cells mediate the onset of thrombosis in the ferric chloride murine model. Blood. 2013;121:3733–3741.

10. Cicilano JC, Sakurai Y, Myers DH, et al. Resolving the multifaceted mechanisms of the ferric chloride thrombsoiss model using an interdisciplinary microfluidic approach. Blood. 2015 126;817–824.

11. Neeves KB. Physiochemical artifacts in FeCl_3_ thrombosis models. Blood. 2015 126;2015:700–701.

