## Supplemental Analysis Guide for "Single-Platelet Mapping of Jugular, Puncture-Wound Thrombi Reveals the Spatial Evolution of Platelet Activation"

**Neighbor analysis**

1. In iVision, use the ‘Measure Segments’ command on a labeled image to generate a text file containing XY coordinates and corresponding segment colors of the platelets.
2. ‘Save As’ a CSV file and open it from Excel.
3. In Excel, copy cells E1:V6 from an existing analysis file and paste.
4. Change the highlighted value (see below) to reflect the # of platelets in cells E2-J2.

*(Same expression is repeated in F2-J2 except for the segment color. Copy up to the last highlighted cell in F2 and paste into G2-J2 as appropriate.)*

5. Highlight cells E2 to J2, double click the bottom right corner to autofill all the data rows.
6. Save as an Excel file.

Description of Excel cells:

Column D: segment color, S1 (5 blue), S2 (4 cyan), S3 (10 green), S4 (2 yellow), S5 (1 red)

Column E: Total neighbors for the platelet found at the XY coordinate in each row.

=SUMPRODUCT(--(SQRT(($B$2:$B$13592-$B2)^2+($C$2:$C$13592-$C2)^2)<$K$1))-1

*(highlighted number is the last row of data, total # of platelet+1)*

Column F-J: Neighbors that are S1 – S5, respectively.

=SUMPRODUCT(--(SQRT(($B$2:$B$13592-$B2)^2+($C$2:$C$13592-$C2)^2)<$K$1),--($D$2:$D$13592=5))-IF($D2=5,1,0)

Cell K1: Neighbor radius in pixels (~ size of a platelet)

L1: Total number of platelets

M1 – Q1: Number of platelets that are S1 – S5, respectively

L2 – L6: Average total number of neighbors for platelets that are S1 – S5, respectively.

M2 – Q2: Average number of neighbors that are S1 – S5 when a platelet is S1.

M3 – Q3: Average number of neighbors that are S1 – S5 when a platelet is S2.

M4 – Q4: Average number of neighbors that are S1 – S5 when a platelet is S3.

M5 – Q5: Average number of neighbors that are S1 – S5 when a platelet is S4.

M6 – Q6: Average number of neighbors that are S1 – S5 when a platelet is S5.

R1 – V1: % of platelets that are S1 – S5, respectively.

R2 – V2: % of neighbors that are S1 – S5, respectively, when a platelet is S1.

R3 – V3: % of neighbors that are S1 – S5, respectively, when a platelet is S2.

R4 – V4: % of neighbors that are S1 – S5, respectively, when a platelet is S3.

R5 – V5: % of neighbors that are S1 – S5, respectively, when a platelet is S4.

R6 – V6: % of neighbors that are S1 – S5, respectively, when a platelet is S5.
